# Assessing the Dose-Dependent Effects of tDCS on Neurometabolites using Magnetic Resonance Spectroscopy

**DOI:** 10.1101/2023.06.13.544864

**Authors:** Anant Shinde, Rajakumar Nagarajan, Muhammed Enes Gunduz, Paul Visintainer, Gottfried Schlaug

## Abstract

Concurrent transcranial direct current stimulation (tDCS) and proton Magnetic Resonance Spectroscopy (^1^H MRS) experiments have shown up- or downregulation of neurotransmitter concentration. However, effects have been modest applying mostly lower current doses and not all studies found significant effects. Dose of stimulation might be an important variable in eliciting a consistent response. To investigate dose effects of tDCS on neurometabolites, we placed an electrode over the left supraorbital region (with a return electrode over the right mastoid bone) and utilized an MRS voxel (3x3x3cm) that was centered over the anterior cingulate/inferior mesial prefrontal region which is in the path of the current distribution. We conducted 5 epochs of acquisition, each one with a 9:18min acquisition time, and applied tDCS in the third epoch. We observed significant dose and polarity dependent modulation of GABA and to a lesser degree of Glutamine/Glutamate (GLX) with the highest and reliable changes seen with the highest current dose, 5mA (current density 0.39 mA/cm^2^), during and after the stimulation epoch compared with pre-stimulation baselines. The strong effect on GABA concentration (achieving a mean change of 63% from baseline, more than twice as much as reported with lower doses of stimulation) establishes tDCS-dose as an important parameter in eliciting a regional brain engagement and response. Furthermore, our experimental design in examining tDCS parameters and effects using shorter epochs of acquisitions might constitute a framework to explore the tDCS parameter space further and establish measures of regional engagement by non-invasive brain-stimulation.

## Introduction

Transcranial direct current stimulation (tDCS) is a non-invasive brain stimulation method capable of modulating neuronal activity in targeted brain regions (Nitsche & Paulus, 2000, 2001; Woods et al., 2016). tDCS is typically applied using two or more electrodes placed over the scalp with an electrical current passing between these electrodes through brain tissue. This electrical current has been shown to have a variety of effects on the brain, including changes in cortical excitability, synaptic neuroplasticity, and modulation of behavioral and cognitive function (Fritsch et al., 2010; Lerud et al., 2021; Nagarajan et al., 2021; Nitsche & Paulus, 2000; A. B. Shinde et al., 2021). The concurrent tDCS-MR setup provides an ideal experimental paradigm in which to examine parameters of stimulation and their neural basis. Magnetic resonance spectroscopy (MRS) allows the measurement of neurotransmitters and metabolites in the brain, such as Gamma-aminobutyric-acid (GABA) (a major inhibitory neurotransmitter in the brain), Glutamate/Glutamine (GLX) (major excitatory neurotransmitters in the brain and commonly lumped together as GLX at 3Tesla MRI strength) as well as other neurometabolites such as N-acetylaspartate (NAA), creatine (Cr), and choline (Cho). MRS can provide information about the underlying biochemical and metabolic changes of non-invasive brain-stimulation in particular its excitatory and inhibitory effects (Amadi et al., 2015; Hunter et al., 2015a, 2015b; Stagg, Bachtiar, et al., 2011c, 2011a; Stagg, Bestmann, et al., 2011; Stagg, O’shea, et al., 2009). However, concurrent tDCS-MRS studies have had variable results which might be related to long acquisition times, subject movements during the long acquisition times, the relatively low dose applied (most studies have used 1-2mA), and the field strength of the MRI machine. Many of these studies have found decreases in GABA following the anodal stimulation on the order of 10-30% from baseline (Antonenko et al., 2017; Bachtiar et al., 2015; Heise et al., 2014; Kim et al., 2014; Patel et al., 2019; Stagg, Bachtiar, et al., 2011b; Stagg, Best, et al., 2009; Tremblay et al., 2013; Wilson et al., 2018). However, a few studies found that anodal tDCS had no effect on GABA (Nwaroh et al., 2020; Wilke et al., 2017).

Most tDCS-MRS studies have focused on applying stimulation to the motor region, specifically the precentral gyrus, while much fewer studies examined neurometabolite effects in non-motor brain regions (Kim et al., 2014; Patel et al., 2019)). The prefrontal lobe is traditionally thought to be challenging for MRS studies (DeMayo et al., 2023). The inferior prefrontal brain regions might exhibit more susceptibility-induced magnetic field distortions which can arise from the presence of air/tissue and tissue/bone interfaces such as around the sinuses and closeness of the skull/brain interface.

Nevertheless, the modulation of GABAergic neurotransmission and GABA concentration in several key regions of the brain has been found to play a major role in several neurologic and psychiatric disorders among them depression when the activity in the inferior cingulate/subcallosal cingulate region is modulated by stimulation, but also in task learning such as motor tasks when the activity is modulated in motor regions. In particular, the intrinsic activity and modulation of activity in the anterior cingulate/inferior mesial prefrontal region has consistently emerged as a core component of major depression pathophysiology and as a marker for monitoring treatment effects (Kito et al., 2012; Mayberg, 2003; Mayberg et al., 1999; Salerian & Altar, 2012; Sankar et al., 2020; Tsolaki et al., 2021). This region is a major focus of directly or indirectly targeted brain-stimulation therapy; indirectly through stimulation of dorsolateral prefrontal cortex (dlPFC) assuming a multisynaptic pathway mediating the effects between dorsolateral prefrontal cortex and the anterior cingulate/inferior mesial prefrontal region or directly through invasively placing deep brain electrodes into the subcallosal cingulate region as a target region.

Our primary objective was to investigate the impact of tDCS dosage on neurometabolites, with a specific focus on GABA, using a novel experimental approach that involved shorter, but repeated acquisition times. This methodology allowed us to explore the reliability of test results and the duration of effects. We chose to target the anterior cingulate/subgenual cingulate region due to its role as a direct or indirect target region of noninvasive and invasive brain stimulation in neuropsychiatric disorders (Mayberg, 2003; Mayberg et al., 1999).

## Methods

### Participants

A total of six participants were recruited from the greater Amherst (MA) area (Females= 4; mean age=26.2 years), each subject participating in multiple sessions. Participants were scanned on a 3T whole-body MR scanner (Magnetom, Skyra, Siemens Medical Solutions, Erlangen, Germany) using a 32-channel head receive coil. All participants were right-handed according to the Edinburgh Handedness Inventory (Oldfield, 1971), had no history of neurologic or psychiatric conditions, and had no contraindications to undergoing MRI and tDCS as verified by a safety checklist. Before the actual concurrent tDCS-MRS sessions, all participants underwent a mock session. The purpose of this mock session was to familiarize them with the experimental setup and ensure their comfort within the magnetic resonance (MR) scanner environment. Additionally, a brief test stimulation was administered during this session to ensure that the participants tolerated the high dose stimulation. This preliminary step helped to ensure that participants were well prepared and at ease for the subsequent tDCS-MRS sessions following several days later. Participants were assigned to one of five concurrent tDCS-MRS sessions on their first day of the experiment. Not all participants were able to participate in all sessions.

### Electrode placement

MR compatible rubber electrodes were placed on each participant’s scalp using the 10-20 Electroencephalogram (EEG) system as a guide for target location. Simulations of current distribution in the brain were performed using SimNIBS software to identify target electrode locations that would achieve highest possible current density in the anterior cingulate/subcallosal cingulate cortex (SCC) region. Magnitude of current density JNorm was measured in the SCC region using three electrode montages. Three electrode montages had target electrode location pairs 1) FP1-C2, 2) F9-C1, and 3) FP1-Mastoid, identified with 10-20 EEG system.

To perform simulation with SimNIBS (Puonti et al., 2020; Saturnino et al., 2019) a high-resolution standard T1 image in MNI space was used. A *unified segmentation* routine implemented in SimNIBS was used for <u>skull</u> segmentation which combines <u>spatial normalization</u>, intensity inhomogeneity correction, and tissue segmentation into one model. The segmentation routine uses a Gaussian mixture model for modeling tissues using the spatial tissue probability map from SPM12. The distribution of current density in the MNI bain was then simulated using the SimNIBS program. The simulation parameter that we are using to determine optimal engagement of the anterior cingulate/SCC is the average **JNorm** of left and right anterior cingulate/SCC regions. **JNorm** is the magnitude of the current density.

The supraorbital region on the left (FP1) was use as a target location while a return electrode was placed over the contralateral mastoid region. The return electrode was 4cm in diameter and the target electrode placed over FP1 was 5cm in diameter. A larger size diameter electrode was used on the exposed skin of the forehead (FP1) to decrease the subjective sensations of tingling and warming, which were more intense at a higher dose. Figure 1 shows a electrode placement for the FP1-contralateral Mastoid montage.

**Figure 1:**
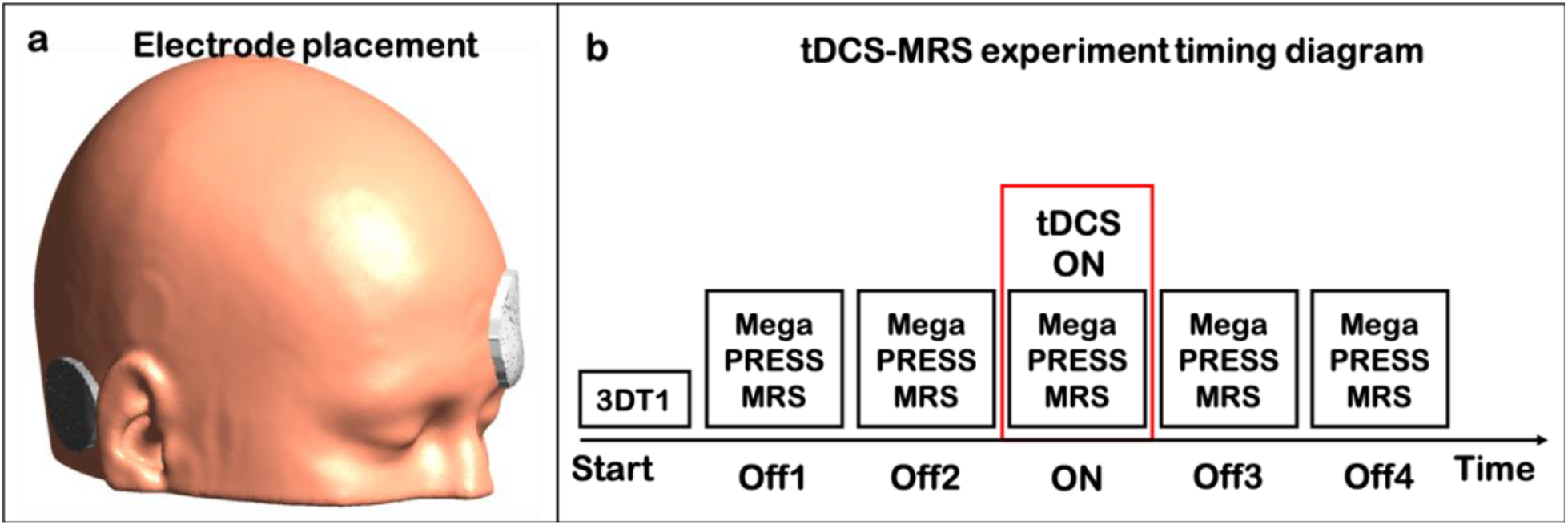
Exeperimental design

### Concurrent tDCS-MRS experiment

After the electrodes were mounted, participants were placed inside the MRI bore in a supine position. Electrodes were connected to the Neuroconn DCMC stimulator device (Neurocare Group, Germany) placed in the MR Control room through a series of connections which ensured that stimulation and MR imaging was not affected by RF noise interference(A. Shinde et al., 2023; A. B. Shinde et al., 2021).

After a participant was situated in the scanner and all connections were secured, a short T1-weighted 3D MPRAGE sequence (resolution= 1.0 × 1.0 × 2.0*mm*^3^; TR/TE 1490/3.36ms; flip angle = 9°; matrix = 256 × 256; field of view = 256 × 256*mm*^2^) with sagittal acquisition and 2mm slice thickness was obtained. This 3D anatomical image set was used to confirm the accuracy of the electrode location with underlying brain regions that we intended to target. If the electrodes were placed inaccurately (meaning that they did not overlay with the target region, confirmed by neuroanatomically experienced investigators), then the participant was moved out of the MR bore, the electrode position was adjusted, and the corrected placement was again confirmed with a fast T1-weighted acquisition. Following the accurate placement of the electrodes, MEGA-PRESS spectra were acquired from a 3 × 3 × 3 cm^3^ voxel centered over the anterior cingulate/subcallosal portions including left and right hemispheres(Nagarajan et al., 2022). An example voxel placement is shown in Figure 2. MEGA-PRESS data were acquired with the following parameters: TR/TE = 2000/ 68ms; 128 averages; 1024 data points; spectral width = 2000 Hz; editing pulse frequencies set to 1.9 ppm and 7.5 ppm for editing of GABA. Water-unsuppressed MEGA-PRESS data were acquired with one average for optimal co-localization of the unsuppressed water signal. MEGA-PRESS sequence was selected as it reliably records GABA and GLX. The complete spectroscopy protocol was around 47 minutes, with a total of five MRS acquisitions of 9 minutes and 18 seconds each. tDCS stimulation was concurrently applied during the third MRS epoch flanked by two non-stimulation scans on either side of the ON condition (OFF1, OFF2, ON, OFF3, and OFF4). Concurrent tDCS-MRS sessions within one subject were separated by 48 hours or more. In each session, a different stimulation was applied with electrode polarity at the mastoid area being either no-stimulation, Anodal 2.5 mA, Anodal 5 mA, Cathodal 2.5 mA, or Cathodal 5 mA. In the no-stimulation session, electrodes were mounted at their respected places, but no current was injected. Participants were not told if and when stimulation was going to occur during the experiment. Also, the session order was varied to avoid any systematic oder effects.

**Figure 2:**
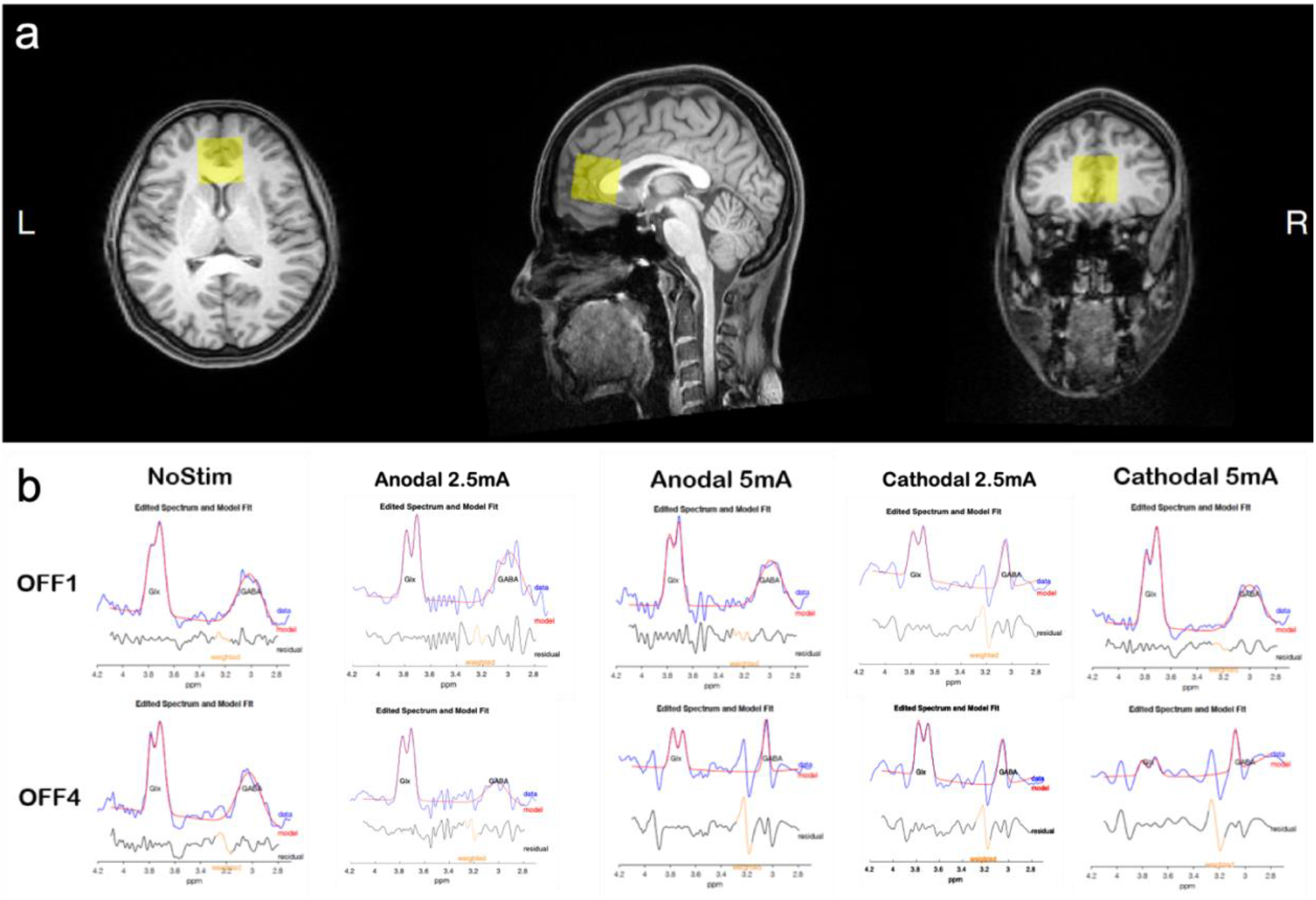
a) MRS voxel placement and b) representative MRS spectrum from at two time poins Off1 and Off4 for each stimmulation dose.

The tDCS stimulation device was programmed to align exactly with the duration of a single MRS acquisition. Stimulation started with ramp up from 0mA to the desired stimulation dose level in a 30 seconds period. The dose level was maintained constant for the duration of the stimulation. Dose was ramped down to 0mA in the last 30 seconds of the stimulation epoch. Timing diagram for the concurrent tDCS-MRS experiment is shown in **Figure 1**.

### Processing

MEGA PRESS sequence generated individual files representing water suppressed and non-suppressed MRS recordings. MEGA-PRESS data were processed using a 5-step process (Load, Fit, Co-Register, Segment, and Quantify) using the MATLAB-based toolbox Gannet 3.2 (Edden et al., 2014; Mikkelsen et al., 2018, 2020). Gannet *‘Load and Fit’* accepts the data in different data formats and maps it to the reference spectra. Gannet preinitialise function helps align acquired spectra with a reference spectrum based on the alignment option. Examples of fitted and aligned spectra for baseline and post stimulation are shown in Figure 2. We used an alignment option called *‘RobustSpecReg*.*’* The Gannet *‘Co-register’* and Gannet *‘Segment’* modules call SPM12 to determine the tissue volume fractions of GM, WM, and CSF. Gannet *‘Quantify’* provides GABA and Glx neurometabolites concentrations in the MRS voxel as well as the fit error against the model function used to calculate these concentrations.

### Statistical Analysis

We used a mixed effects model that accounts for repeated measures and missing data to evaluate effects on GABA and GLX. Models for GABA and GLX were developed separately. Each model has two fixed effects namely **Dose** (-5mA, -2.5m, 0mA, 2.5mA, and 5mA) and **Time of Measurement** (OFF1, OFF2, ON, OFF3, and OFF4). Dose of 0mA or No-stim was considered as a baseline and epoch OFF1 was considered as baseline. We evaluated a dose-by-time interaction to determine whether responses by dose differed over the time measurement. Since the interaction terms were not significant in either model, they were dropped from the analyses. A global test of main effects were conducted using likelihood ratio or Wald tests. If the global test was significant, post-hoc analysis were conducted using Sidak’s adjustment for multiple comparisons. Tolerability scores were analyzed using the Kruskal-Wallis non-parametric approach. Post-hoc tests were conducted Bonferroni’s adjustment.

### Safety and tolerability

After each session of tDCS, we recorded safety and tolerability information. The skin and scalp locations under the electrodes were inspected for any skin burns or other lesions. At the end of each session, we asked volunteers to indicate their tolerance of the noninvasive <u>brain stimulation</u> on a <u>visual analog scale</u> (VAS) with 0 and 10 as the endpoints where zero indicated that subjects tolerated the stimulation well and had no unusual sensations and 10 indicated that the stimulation session caused strong sensory experiences and was judged to be barely tolerable. Since the range of tolerability scores was limited and sample size was small, a Kruskal-Wallis test was run to analyze data.

## Results

The simulation parameter that we are using to determine optimal engagement of the anterior cingulate/SCC is the average **JNorm** of left and right SCC region. **JNorm** is the magnitude of the current density that can be detected in the anterior cingulate/subcallosal cinguate regions. The figure below (**Figure 3**) shows three montages that we experimented with. The montage with the electrode on FP1 and the return electrode over the contralateral mastoid gave us the highest **JNorm** with values at least 20% higher than the next best montage; this montage was used to evaluate effects of different stimulation doses on the neurometabolites in the SCC region.

**Figure 3:**
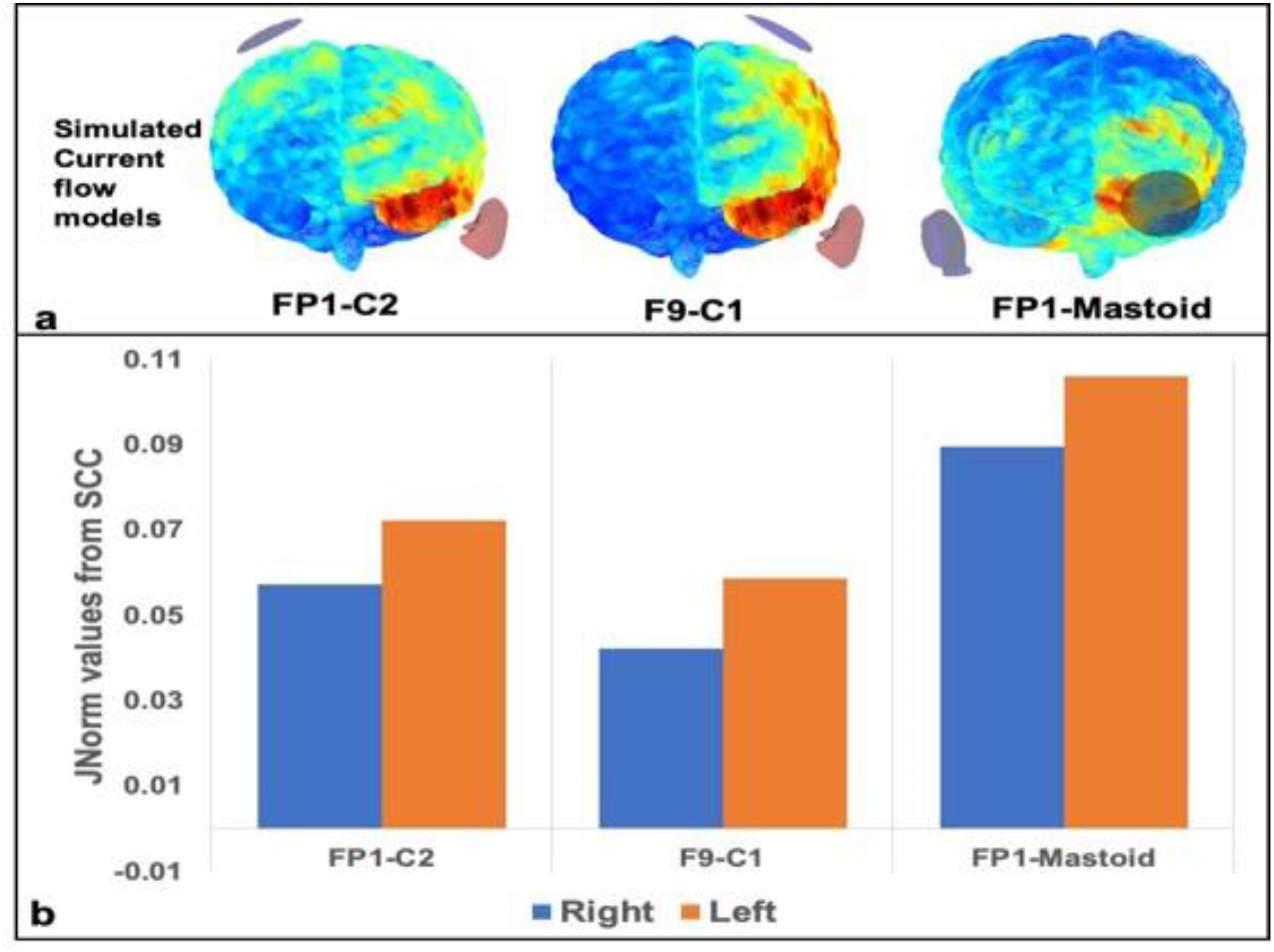
Three different montages are presented (6a – frontal view) with their respective JNorm values (6b). The electrodes over the FP region and the contralateral mastoid region achieved the highest JNorm values (6b) in a Subcallosal Cingulate (SCC) region.

Figure 4 shows the mean changes in GABA and GLX neurometabolites relative to the OFF1 reference/baseline. GABA values in the selected MRS voxel showed large changes with 5mA applied while changes in GABA was much smaller when the 2.5mA dose was applied. Polarity had an opposite effect on the GABA response. The GABA values returned closer to the baseline during the second MRS after the stimulation. GLX values did not show large changes similar to GABA. However, consistent downregulation post-stimulation was observed in GLX when cathodal 5mA tDCS was applied compared to No-Stim, the downregulation was also observed at timepoints Off3 and Off4 compared to Off1 baseline. Further, the effects seemed to be slightly more delayed for GLX than they were for GABA; changes in GLX concentration were more evident post stimulation in contrast changes in GABA were immediate which were observed in the MRS acquired while stimulation was applied.

**Figure 4:**
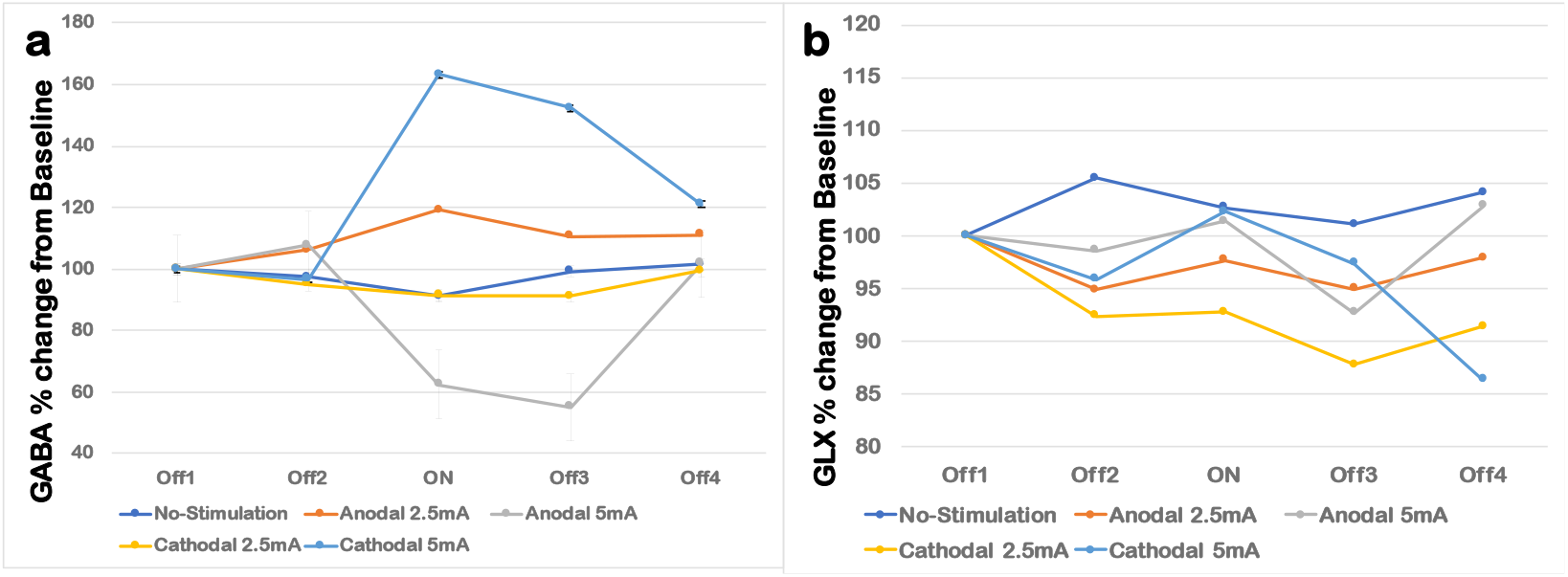
a)% Change in GABA at different timepoints. B) % Change in GLX at different timepoints. [different scales are chosen for better representation of changes]

### Mixed Effects analysis of GABA values

The overall marginal model achieved a significant result with a p-value of 0.0013. Time factor had no significant effect on GABA values, i.e. no significant change in GABA value was observed at any timepoint when compared to Off1. This could be attributed to the polarity dependent changes in GABA values which increased with cathodal stimulation and decreased with anodal stimulation. Specifically, Anodal 5mA dose showed significant change in GABA compared to No-Stim baseline (p =0.001). There was no significant effect of interaction between stimulation and time. Additionally, a post-hoc analysis comparing stimulation categories, adjusting for multiple comparisons using Sidak’s method, showed significant change in GABA concentration with three stimulation pairs: 1) No-stim vs. Anodal 5mA (p=0.008), 2) Anodal 5mA vs. Anodal 2.5mA (p=0.016), and 3) Cathodal 5mA vs. Anodal 5mA(p=0.000).

### Mixed Effects analysis of GLX values

Overall marginal model did not reach statistical significance (p = 0.1697). However, Timepoint Off3 immediately after ON significantly differed in GLX values compared to baseline (p=0.048). Anodal 5mA caused a significant decrease in GLX concentration compared to No-Stim baseline (p =0.031). There was no significant effect of interaction between stimulation dose and timepoint.

### Tolerability Scores

High dose tDCS stimulation did not lead to any adverse skin or neurological effects among the participants. None of the participants experienced severe headaches, seizures, neurological impairments, or skin burns.

In addition, stimulation was well tolerated by all the participants. However, none of the participants stopped the experiment due to the stimulation being intolerable (which we defined as a VAS score of 9 and 10).

Figure 5 shows the mean tolerability scores for each stimulation dose. It is evident that higher dose correspond to higher but tolerable VAS scores. The overall K-W test is significant (p = 0.014, adjusting for ties). Post-hoc analysis showed significant difference(p=0.001296) in No stimulation and Cathodal 5mA group after Bonferroni correction.

**Figure 5:**
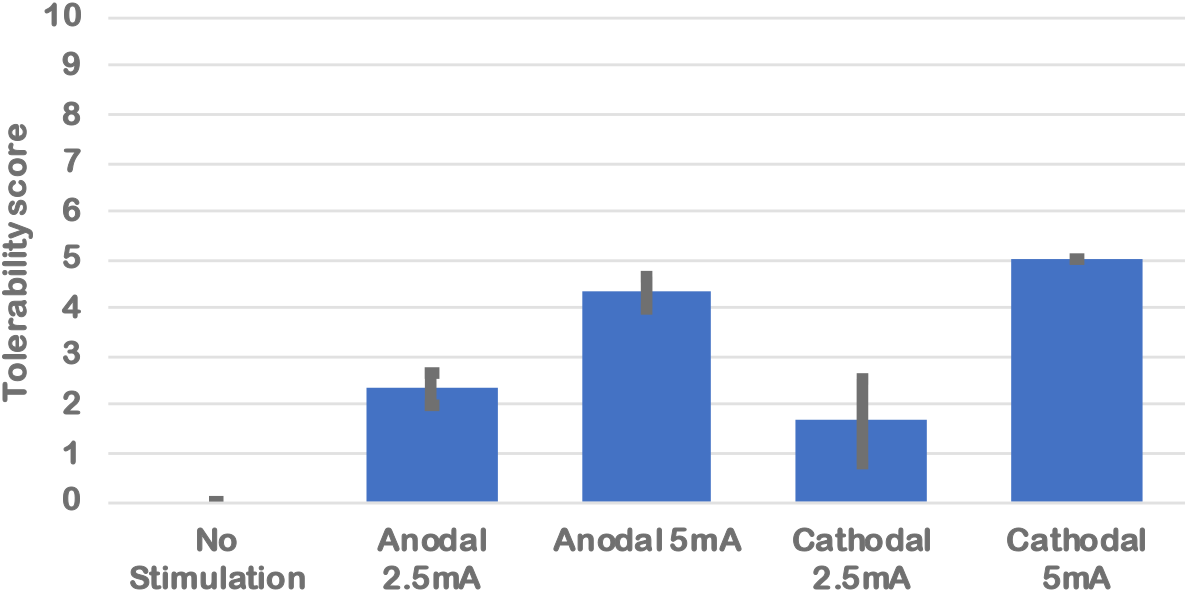
Mean tolerability scores vs stimulation (mean +/-SEM)

## Discussion and Conclusions

Simulations and experiements conducted in this study suggest that high dose tDCS can reliably modulate neurometabolite concentrations such as GABA and GLX in an anterior cingulate/inferior mesial prefrontal region of the brain showing a clear dose and polarity effect. The highest dose of 5mA caused strong and consistent effects with an average of 63% change in regional GABA concentrations. Lower dose 2.5mA stimulation showed a change of approximately 20%. There was minimal change in GLX concentrations when the stimulation was applied with change from baseline being consistently lower than 20% at all timepoints. Interestingly, immediate post-stimulation MRS measurements demonstrated a decrease in GLX compared to baseline, suggesting that the impact of stimulation on GLX at this specific time point is independent of stimulation polarity.

Previous studies have provided evidence that the excitatory effects of anodal tDCS are associated with a modulation of GABAergic interneurons (Antal et al., 2004; Liebetanz et al., 2003; Nitsche et al., 2004; Stagg, Best, et al., 2009; Stagg, Wylezinska, et al., 2009) which might be due to down regulation of the enzyme GAD-67 activity (Lau & Murthy, 2012; Sahyoun et al., 2004) showed that anodal tDCS lead to a decrease in GABA values during and after stimulation of motor region but did not affect Glutamate concentration(Bachtiar et al., 2018). Reduction in GABA values was also observed in other studies that applied anodal stimulation to the visual cortex of the cats and inferior frontal gyrus of the humans(Harris et al., 2019; Zhao et al., 2020). A few a-tDCS studies have shown no modulation of GABA concentration or GABA receptor activity (Dwyer et al., 2019; Nwaroh et al., 2020; Wilke et al., 2017), most likely because of lower dose of stimulation and the necessities of long acquisition times. Most studies have evaluated effects of stimulation (doses between 1 to 2 mA applied for 10 to 30 minutes) on the GABA values in the cortical regions underneath the tDCS electrode. Our findings regarding the changes in GABA levels following anodal tDCS align with previous research. Moreover, we observed that higher tDCS doses have the potential to induce greater modulation of GABA values (∼63%) compared to lower doses (∼20%) when compared to baseline levels. Changes in GABA values were close to 20% when 2.5mA tDCS dose was applied which is similar to other studies that evaluated effect of tDCS dose of 2mA. Researchers have also shown that cathodal tDCS increases concentration of GABA in the brain region underneath the electrode (Hasan et al., 2012; Heimrath et al., 2020; Zhao et al., 2020). Our results coroborate these with observed increases in GABA values when cathodal tDCS was applied.

Given that GABAergic cortical inhibitory interneurons play a role in several neurological disorders such as Anxiety, Depression, Epilepsy, Alzheimer’s disease (Koliatsos et al., 2006; Pham et al., 2009; Sarawagi et al., 2021), modulation of these GABA interneurons by tDCS could be a potential disease/disorder-modifying mechanism for improving function. The use of high-dose tDCS has the potential to induce stronger and more reliable effects, making it a promising candidate as a facilitator in the treatment of various neurological and psychiatric disorders. By applying higher doses of tDCS, we can enhance the modulation of neurometabolites, such as GABA. This could potentially augment the therapeutic potentials of tDCS, either as a standalone treatment or in combination with other existing medical therapies. Intrinsic activity and modulation of activity in the anterior cinculate and specifically in the subcallosal cingulate (SCC) subregion has consistently emerged as a core component of major depression pathophysiology(Mayberg, 2003; Mayberg et al., 1999) and as a marker for monitoring treatment effects (Kito et al., 2012; Salerian & Altar, 2012; Sankar et al., 2020; Tsolaki et al., 2021). The anterior cingulate/inferior mesial prefrontal region stimulation of dorsolateral prefrontal cortex (dlPFC) assuming a multisynaptic pathway mediating the effects between dorsolateral prefrontal cortex and the SCC region or directly through invasively placing deep brain electrodes into the SCC region as a target region. Our results and concurrent tDCS-MRS approach provide an experimental framework to show the engagement of this region through non-invasive brain-stimulation and the modulations of major neurotransmitters of this region, which can be further explored in several neurologic and psychiatric disorders.

Previous in-vivo human studies have reported inconsistent and small modulations of the neurometabolite GLX with anodal tDCS stimulation (Harris et al., 2019b; Heimrath et al., 2014, 2020; Zhao et al., 2020b). In our study, we observed small but reliable changes in GLX during and after stimulation in the anterior cingulate/inferior mesial prefrontal region. These findings suggest that the effects of tDCS on GLX may vary depending on the brain region being stimulated. Further research is needed to replicate these findings and investigate the effects of high-dose stimulation on GLX in other brain regions.

Our experimental design, which incorporates shorter image acquisition and stimulation times, presents a novel framework for investigating the biological effects of various stimulation parameters. By utilizing shorter acquisition times, we gain greater flexibility in testing different parameters such as stimulation dose, polarity, electrode montage, and duration. This design also allows us to compare the real effects of stimulation with the test-retest variations observed in the absence of stimulation (no-stimulation condition). Additionally, by examining the duration of induced effects, we can assess the persistence and temporal dynamics of the observed changes. Furthermore, our design enables us to explore the potential synergistic effects of high-dose brain stimulation in combination with other interventions. Overall, this experimental framework provides a valuable approach for advancing our understanding of the effects of brain stimulation and its potential integration with other therapeutic strategies.

## Acknowledgments

This research was supported by an NIH Brain Initiative grant (**7R01MH111874-05**). GS also acknowledges support from U01NS102353. AS acknowledges support from an UMass-Initiative-on Neurosciences (ION) Inspiration Awards for Neuroscience and Technology. We are thankful for and very much appreciate the generous support that Klaus Schellhorn from neuroConn has provided to us with making a state-of-the-art multi-channel MR DC stimulator available to us and his helpful suggestions with setting up the concurrent tDCS-MR experiments and being available for anytrouble shooting over the years. We would also like to thank Elena Bliss, our MR Technologist, for her support with all of the complicated and lengthy concurrent tDCS-MRS acquisitions.

